# Four-color fluorescence cross-correlation spectroscopy with one laser and one camera

**DOI:** 10.1101/2023.01.30.526256

**Authors:** Sonali A. Gandhi, Matthew A. Sanders, James G. Granneman, Christopher V. Kelly

## Abstract

The diffusion and reorganization of phospholipids and membrane-associated proteins are fundamental for cellular function. Fluorescence cross-correlation spectroscopy (FCCS) measures the diffusion and molecular interactions at nanomolar concentration in biological systems. We have developed a novel, economical method to simultaneously monitor diffusion and oligomerization with the use of super-continuum laser and spectral deconvolution from a single detector. Customizable excitation wavelengths were chosen from the wide-band source and spectral fitting of the emitted light revealed the interactions for up to four spectrally overlapping fluorophores simultaneously. This method was applied to perform four-color FCCS, as demonstrated with polystyrene nanoparticles, lipid vesicles, and membrane-bound molecules. Up to four individually customizable excitation channels were selected from the broad-spectrum fiber laser to excite the diffusers within a diffraction-limited spot. The fluorescence emission passed through a cleanup filter and a dispersive prism prior to being collected by a sCMOS or EMCCD camera with up to 10 kHz frame rates. The emission intensity versus time of each fluorophore was extracted through a linear least-square fitting of each camera frame and temporally correlated via custom software. Auto- and cross-correlation functions enabled the measurement of the diffusion rates and binding partners. We have measured the induced aggregation of nanobeads and lipid vesicles in solution upon increasing the buffer salinity. Because of the adaptability of investigating four fluorophores simultaneously with a cost-effective method, this technique will have wide application for examining complex homo- and heterooligomerization in model and living systems.

## 1. Introduction

Fluorescence correlation spectroscopy (FCS) and fluorescence cross-correlation spectroscopy (FCCS) are powerful techniques to reveal the diffusion behavior of molecules at nanomolar concentration in model and biological systems (1–7). FCS and FCCS provide single-molecule sensitivity to reveal averaged diffusive characteristics through a diffraction-limited confocal volume (8–10). As fluorescently labelled molecules diffuse through a detection volume, the fluctuating fluorescent signals are obtained and correlated (11, 12). The fluorescence emission intensity versus time varies depending on the number and brightness of the fluorophores as they enter and leave the observation volume (13–16). FCS reveals biophysical parameters of a single-color channel, including the molecular diffusion coefficients and concentrations by analyzing the autocorrelation dwell time (*τ*_*D*_) and amplitude (*G_0_*). The comparison of multiple color channels occurs with fluorescence cross-correlation spectroscopy (FCCS) (17) during which the molecular interactions and co-diffusion are measured (2, 11). If distinctly labelled particles are bound together as they diffuse through the observation spot, then their fluorescence intensities have simultaneous fluctuations and their cross-correlation amplitude is large (18).

FCS and FCCS typically examine fluorescence emission with an acquisition rate of ≤10 kHz and <10 μW of excitation light focused to a diffraction-limited observation spot. The fluorophores are excited and emit thousands of times as they diffuse through the observation volume. For this purpose, a sensitive detector and well-chosen fluorophores with complementary excitation and emission spectrums are prerequisites. Fluorophores with high absorption cross-sections, high quantum efficiencies, and resistance to photobleaching are desired to obtain 10-1000 collected photons during their microsecond transit through the detection volume (3).

To observe molecular binding with FCS the bound and unbound states must typically have diffusion coefficients that vary by 10x, which is rare upon ligand binding or dimerization (19, 20). FCCS overcomes this limitation by detecting free diffusors and aggregates through multi-color correlations (21, 22). For FCCS, fluorescent molecules are labelled with chromatically distinct fluorophores and the cross-correlation of the corresponding detection channels reveals their co-diffusion (21, 23, 24). Generally, the emission of all diffusers is obtained simultaneously that have same experimental conditions (2).

Traditional, FCCS is limited to the observation of various fluorophores that are well chromatically separated such that minimal fluorescence emission bleed-through is detected in the wrong channel (25, 26). Typically, the color channels are chromatically separated through dichroic mirrors and channel-specific emission filters prior to each channel’s detection in a dedicated avalanche photodiode or photomultiplier tube. Bleed-through between the color channels often yields a non-zero cross-correlation for independently diffusing fluorophores and difficulty in observing molecular binding events. Crosstalk between color channels is especially problematic when fluorescent proteins are used, and each fluorophore’s emission spectrum is wide.

To overcome this crucial spectral cross-talk, pulsed interleaved excitation (PIE) FCCS method has been developed where excitation and detection of each fluorophores occurs at unique time points (17, 27). In addition to separating the emission spectrally as in standard multicolor FCCS, temporal separation via picosecond pulsed excitation single photon emission counting provides greater fluorophore resolution (28). However, this technique often requires numerous fluorophore specific detectors, and multiple pinhole alignments for each emission channel. Also, previous studies on prism-based FCCS simplify the detection path of the FCCS experiment by using a dispersive prism to spectrally separate the emission light distinctly for spectral ranges (2, 23, 29), but it have had high background noise for hyper spectral detection.

This manuscript reports on a new FCCS method that can resolve up to four spectrally overlapping fluorophores to study their mobility and clustered interactions with a single excitation source and a single detector. This method simultaneously measures the intensity versus time of four fluorophores to be analyzed as four autocorrelations, six two-color cross-correlations, and up-to five higher-order cross-correlations. The advantage of multicolored FCCS is that it enables measurement of molecular density and diffusion rates for four populations. In addition, the emission from four channels is also useful to calculate cross-correlation between the pairs to determine oligomerization. This novel method uses a single super-continuum laser as an excitation source that provides multiple excitation wavelengths dependent on the choice of excitation filters. A single confocal aperture and dispersive prism play vital roles by reducing out-of-focus fluorescence emission to enhanced SNR and chromatically spread emission on single sensor that obviate the need of multiple detectors, respectively. The article provides details on the experimental setup and demonstrates its capability to resolve four fluorescent polystyrene nanobeads and three fluorescently labelled lipid vesicles in solution. FCCS revealed that the nanobeads and the lipid vesicles initially diffused independently with negligible cross-correlations. Upon increasing the buffer osmolarity or the addition of bovine serum albumin (BSA), the nanobeads and the lipid vesicles oligomerized into multi-colored diffusers with significant cross-correlations. Further, single-molecule detection has been verified by revealing the independent diffusion of membrane-bound fluorescent proteins and fluorescently labeled phospholipids. Future adaptations of this method will provide resolution and interactions of the intricate binding and unbinding events of membrane-bound proteins.

## 2. Materials and Methods

### 2.1 Sample preparation

Four different single-color fluorescent nanobeads were mixed together and diluted with milli-Q water with a resistivity >18 MΩ cm. 100 nm diameter blue beads with an excitation wavelength (*λ_ex_*) of 450 nm (Fluoro-Max; Fisher Scientific), 100 nm diameter yellow bead; *λ_ex_* = 515 nm (FluoSpheres; Life Technologies), 100 nm diameter red bead; *λ_ex_* = 561 nm (Fluoro-Max; Fisher Scientific), 40 nm diameter dark red bead; *λ_ex_* = 635 nm (FluoSpheres, Life Technologies). The mixture of nanoparticles was placed on a glass bottom dish (MatTek) after the dish was rinsed with ethanol and dried in nitrogen stream for 15 sec.

Large unilamellar vesicles (LUVs) were composed of non-fluorescent phospholipids and fluorescent phospholipids. Four different lipid mixtures were used 1,2-dioleoyl-sn-glycero-3-phosphocholine (DOPC; Avanti Polar Lipids), 1-palmitoyl-2-(dipyrrometheneboron difluoride)undecanoyl-sn-glycero-3-phosphocholine (TF-PC, Avanti Polar Lipids), 1,2-dipalmitoyl-sn-glycero-phosphoethanolamine Texas Red (DPPE-TR, Life Technologies), 1,2-dioleoyl-sn-glycero-3-phosphocholine-N-Cyanine 5 (DOPC-Cy5, Avanti Polar Lipids). DOPC (99.98 mol%) and TF-PC, DPPE-TR, or DOPC-Cy5 (0.02 mol%) in chloroform were mixed in glass vial. Lipids were dried under a nitrogen stream and kept in vacuum for >30 min. The lipid film was hydrated to 1 mg/mL with milli-Q water (<18 mΩ). The solution was vortexed and extruded eleven times through membrane filter with a 100 nm pore size within a liposome extruder (LiposoFast, Avestin) to obtain vesicles. The procedure was repeated to yield independent stock solutions for each color LUVs before they were mixed in single vial to examine their independent or correlated diffusion.

Supported lipid bilayers (SLBs) were prepared by the bursting of giant unilamellar vesicles (GUVs) upon a glass coverslip. GUVs of 1-palmitoyl-2-oleoyl-snglycero-3-phosphocholine (POPC; Avanti Polar Lipids) and 0.3 mol % GM1 Ganglioside (Avanti Polar Lipids) were prepared by electro-formation, as described previously (30–33). Exposing the GUVs on plasma-cleaned coverslips resulted in their bursting and created continuous bilayer. CTxB was labeled with AlexaFluor 647 or AlexaFluor 488 before purchase (Thermo Fisher Scientific). CTxB was added to the SLB for a final concentration of 0.25 mg/mL above the SLB to saturate all available GM1. After 0.5 min of incubation, the unbound CTxB was rinsed away.

The phospholipid monolayers of the artificial lipid droplets (aLDs) were formed from multilamellar vesicles (MLVs) composed of non-fluorescent DOPC and 0.05 mol% fluorescent DOPC-Cy5. The phospholipids were mixed in chloroform, dried under a nitrogen stream, kept under vacuum for >30 min. The dried phospholipid film was hydrated to a concentration of 0.1 mg/mL Intracellular Buffer (IB) and lightly vortexed to obtain MLVs. IB consists of 10 mM HEPES, 140 mM KCl, 6 mM NaCl, 1mM MgCl_2_, and 2 mM EGTA within milli-Q water. The MLVs were mixed with glyceryl trioleate (TO; Millipore Sigma) at a mass ratio of 1000:1 TO:PL and bath sonicated for 30 min (Ultrasonic Cleaner 96043-936; VWR Symphony) to form aLDs. aLDs were mixed with 20 μm diameter nonfluorescent polystyrene beads (Millipore Sigma) and purified proteins, as needed, prior to being sandwiched between a glass slide and a coverslip (Fisher Scientific), which were previously cleaned and passivated. Details for our protein purification are given in the supplemental material. The glass slides and coverslips were rinsed with ethanol, dried under nitrogen stream, and passivated with a casein. Passivation included exposing the slides and coverslips to 20 mg/mL of casein in milliQ water for >20 min prior to rinsed again with milli-Q water (34). 7 μL of the aLD, bead, and protein mixture was sandwiched between a glass slide and coverslip as the final stage in sample preparation. In this system, the nonfluorescent beads acted as spacers to prevent the coverslip and the glass slide from getting too close together and smashing the aLDs. The buoyant aLDs floated at the top of the sandwich, which was the glass slide while the coverslip was closer to the inverted microscope objective. The bottom of the aLDs, the “south pole,” was the focus for our FCCS observation point such that neither the illumination nor emission light had to refract through the aLD on its path to or from the objective.

### 2.2 Optical setup

The four-color FCCS was performed with a customized inverted microscope (IX71, Olympus) on a vibration isolated optical table. Excitation was provided by a super-continuum fiber laser (SC-Pro, YSL photonics) that produced broad spectrum pulsed laser with wavelengths ranging from 400 nm to 2400 nm. For safety, the long-wavelength component (>650 nm) was separated by a dichroic mirror and blocked within a light-tight box to reduce the total laser power present during the free-space customization of the fluorescence excitation paths. The optical colors were separated by dichroic mirrors, chromatically filtered (BrightLine FF02-435/40, FF01-513/13, FF01-561/14, and FF01-630/38, Semrock), expanded, attenuated, and recombined. Each excitation color had <10 μW average power. The combined excitation channels were reflected by a three-band dichromic mirror (ZT442/514/561rpc, Chroma) and directed into a 40x, 1.30NA microscope objective (UIS2 BFP1, Olympus) to a focused point in the sample. The emission from the diffraction-limited illumination point was collected by the same objective, passed through the three-band dichroic mirror, transmitted through the emission filter (ZET442/514/561m, Chroma), and focused by the microscope tube lens through a 40 μm diameter confocal pinhole (P40D, ThorLabs). The emission was transmitted through a relay lens (Optomask, Andor) that was customized with the insertion of a chromatically separating prism (PS812-A, ThorLabs). The emission was collected on the center pixels of either an EMCCD (iXon-897 Ultra, Andor) or sCMOS camera (Zyla, Andor) with the emission chromatically spread on the camera sensor (Fig. 1). This setup provided a 16 nm color difference per pixel on the 13 μm wide pixels of the EMCCD. Both the confocal pinhole and the camera were placed on 3D translation stages (PT3, ThorLabs) to allow for precise alignment. This setup has integrated computer control connected with camera via custom LabVIEW routines (National Instruments).

**Figure 1:**
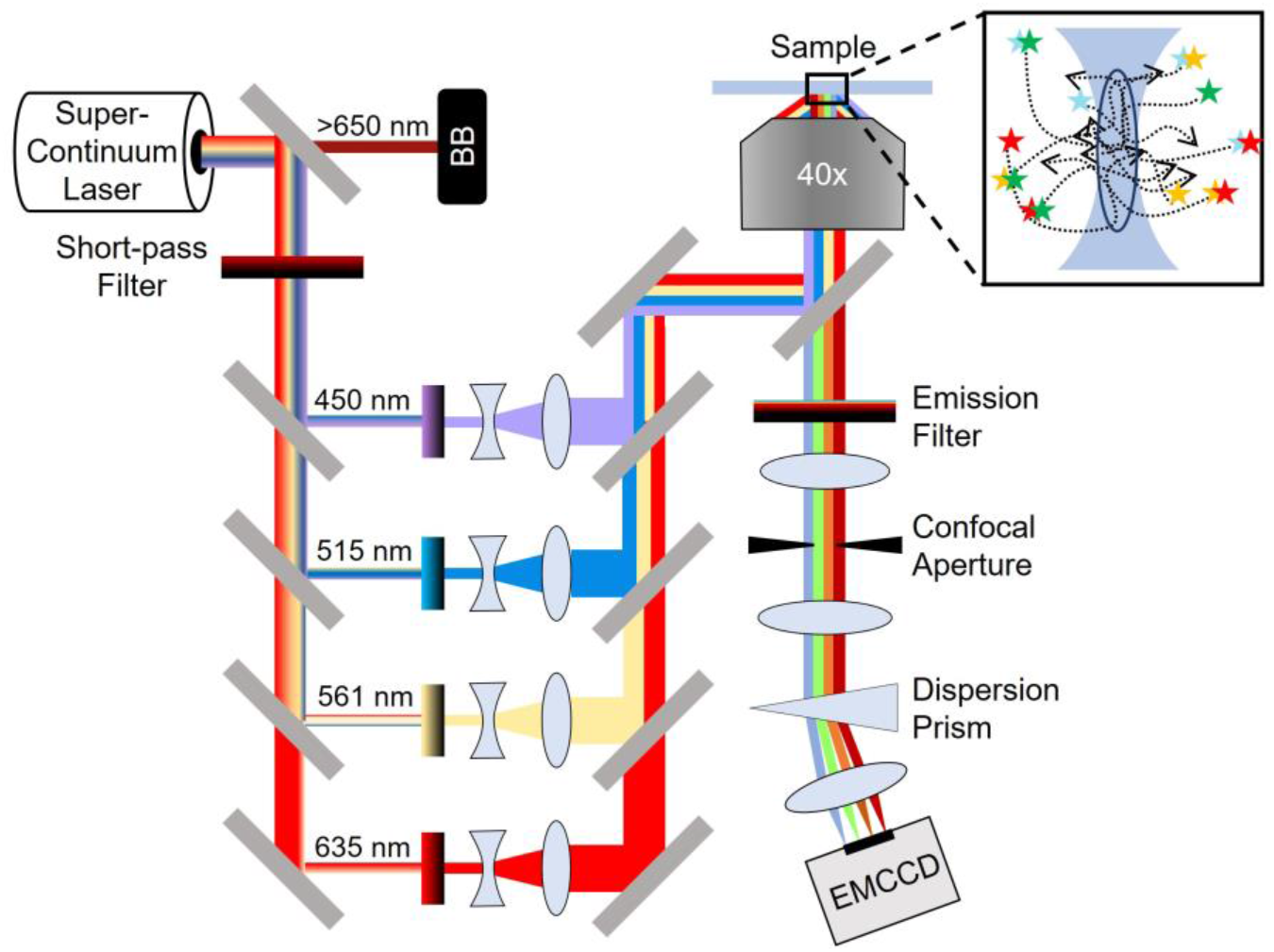
The FCCS set up included a super-continuum laser that was separated into four narrow-spectrum excitation channels that travelled through clean-up filters, neutral density filters, and beam-expanding telescopes prior to being recombined. The combined excitation colors were passed through the microscope objective to illuminate a diffraction-limited volume within the sample. The fluorescence emission was collected by the objective, passed through an emission filter and a pinhole before being focused on the camera sensor. (Inset) Four fluorescently labelled species diffusing independently or as an oligomer through the confocal detection volume yield independent or correlated emission intensities.

The excitation beams required precise alignment and determination of the size of illumination volume. Fluorescent supported lipid bilayer (SLB) was imaged when illuminated by each focused excitation beam. SLBs provided a uniform fluorophore distribution such that the fluorescence image without a pinhole showed the shape of the illumination. The image of the excitation light on the SLB was fit to a 2D Gaussian function to determine the width of laser spot (*w_0_*) (Fig. S1). To confirm the 3D confocal volume, the diffusion of 100 nm diameter fluorescent polystyrene particles was compared to expected results. The diffusion coefficient (*D*) was calculated theoretically, *D* = *k_B_T*/(3*πηd*), and measured experimentally, *D* = *w*_0_^2^/(4*τ*_*D*_), which incorporate Boltzmann’s constant (*k_B_*), the temperature (*T*), solvent viscosity (*η*), particle diameter (*d*), and dwell time (*τ*_*D*_). The experimental and theoretical values of *D* for 100 nm diameter polystyrene microspheres in water were consistent, *D* = 4.45 ± 0.06 μm^2^/s and 4.39 μm^2^/s, respectively (35).

Samples were exposed to <10 μW total excitation power with *λ_ex_* = 450, 515, and 561 nm. Our optical setup also provides a 635 nm excitation, but the fluorophores with >680 nm emission used in these experiments were sufficiently excited by the shorter wavelength excitation such that the 635 nm excitation was not used here. Each EMCCD camera frame was acquired with a 0.1-ms exposure at a 1,785 Hz frame rate. 53,500 frames were acquired per 30-sec acquisitions for the 496×4 cropped region of interest (ROI) with the EMCCD. Similarly, sCMOS acquired with 5 ms exposure time and 196 Hz frame rate that yielded 5876 frames per 30 sec acquisitions for 20×4 cropped ROI. This frame rate from EMCCD is sufficient to measure the diffusion of membrane-bound molecules or >40 nm diameter diffusers in solution for which diffusion rates are typically <10 μm^2^/s. However, the frame rate was not sufficient to measure small molecule diffusion in aqueous suspension for which diffusion rates are typically >100 μm^2^/s.

### 2.3 Extraction of I versus T

Each image was averaged over the dimension perpendicular to the chromatic separation to isolate the emission spectrum from all fluorophores at a moment in time. This spectrum versus time was analyzed to extract the relative changes to each fluorophore’s concentration within the observation spot versus time. Extraction of each fluorophore’s concentration required determining the expected emission spectrum from each fluorophore and fitting each acquired spectrum to a linear combination of the spectra from the present fluorophores.

The fluorescence emission spectrum for each fluorophore was collected from control samples that each contained one fluorophore while excited by all excitation colors (Fig. 2A-D, S3A-C). To account for day-to-day sub-pixel misalignment, the single-fluorophore spectra were each fit to the sum of three Gaussians to provide a continuum model of each fluorophore’s emission after the three-band dichroic mirror and emission filter (Figs. 2F, S3E),

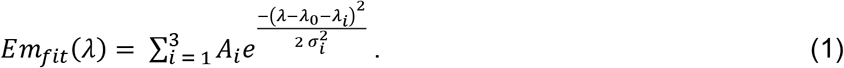

**Figure 2:**
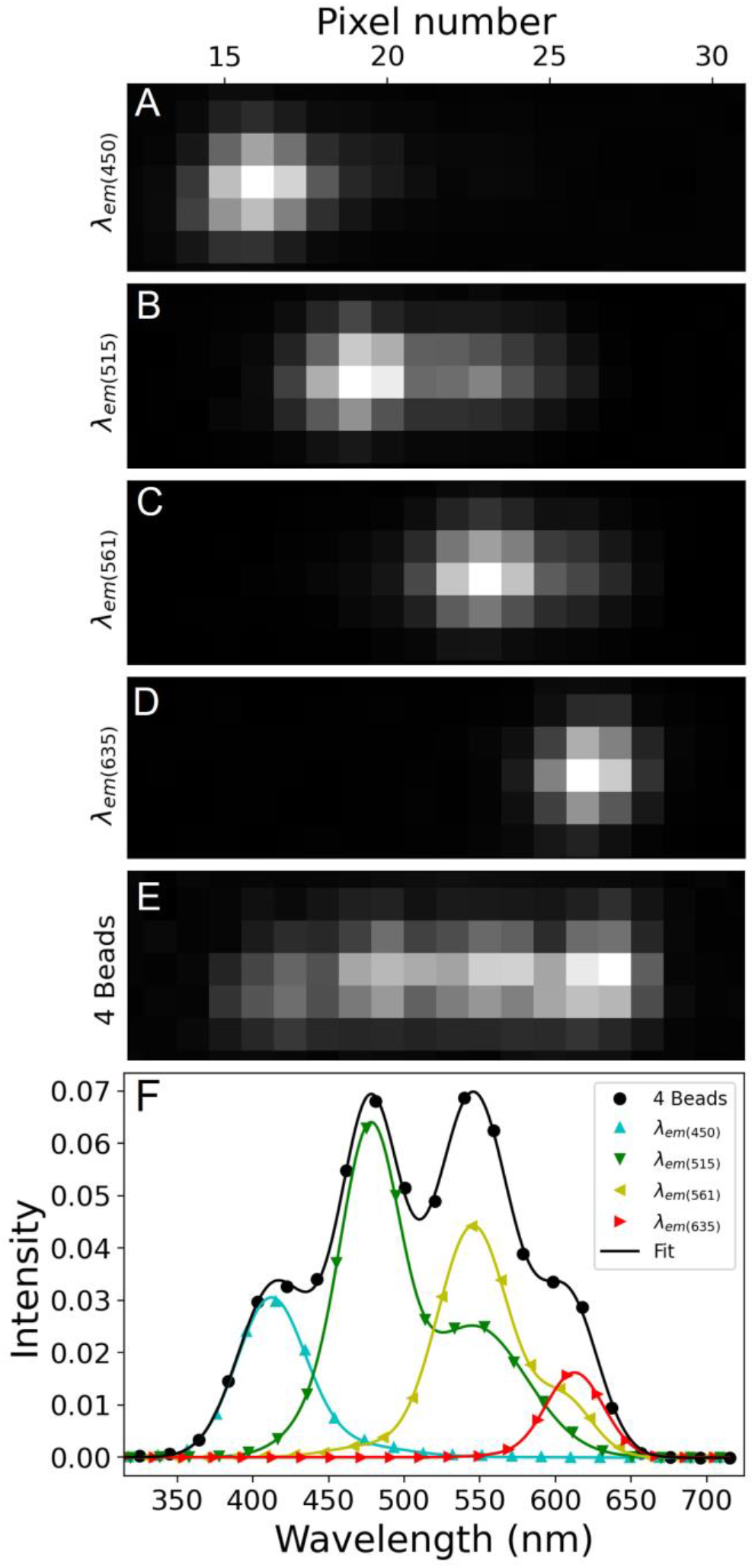
Chromatic separation of emission spectrum on the camera sensor. Spectrally overlapping emission of fluorescent nanobeads was dispersed via prism was collected on the camera. (A-D) Single nanobeads of excitation wavelengths 450, 515, 561 and 635 nm, respectively, were spectrally separated and collected on the cropped ROI. Each column of pixels on the camera is associated with an emission wavelength. (E) Samples with all four nanobeads show the distinct, but highly overlapping emission spectra. (F) Control samples with only one color of bead present were used to identify the emission spectrum of each bead type (*colored symbols and fits)*. With four nanobeads simultaneously present, the spectrum was fit to reveal the emission intensity of each bead type for each camera frame (*black symbols and fit)*.

This fitting function was chosen because it provided a smooth, analytical fit to the collected points of the single-fluorophore emission spectra with fewer fitting parameters than other candidate fit functions; there was not a physical meaning for this three-Gaussian fit. Eq. 1 provided a continuous function that represented the spectrum of each fluorophore used. The nine fit parameters of amplitude (*A_i_*), center (*λ_i_*), and width (σ_i_) were identified for each fluorophore and unchanged during later fits. The fitting parameter *λ_0_* was shared by all three Gaussians and allowed for the emission spectra for each fluorophore to be effectively shifted on the sensor to account for day-to-day sub-pixel variations in optical alignment. Another control sample was created with well-resolved intensities of *M* different fluorophores simultaneously to determine the relative shift of the calibration files with a separate *λ_0_* for each fluorophore (*λ_0,Fluor_*) (Figs. 2E, F,S3D,E) according to

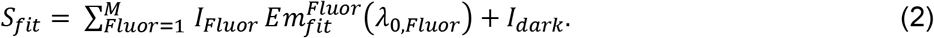

When fitting the sum of each fluorophore’s 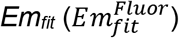 to this multi-colored control sample the relative shift (*λ_0,Fluor_*) and the amplitude (*I_Fluor_*) of each 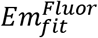 were identified. Later fits kept *λ_0,Fluor_* fixed because *λ_0,Fluor_* represents the inherent relative emission spectrum of each fluorophore. The dark counts on the detector, which were approximately 2% of the signal, were subtracted during fitting by the incorporation of a constant (*I_dark_*).

The system required sample-specific sub-pixel alignment. A shift of the emission spectra was determined for each sample while keeping the relative shift for each fluorophore constant. For each sample, the time-averaged sample spectrum was fit with a single shared shift for all 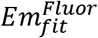 corresponding to the sample-specific chromatic shift on the camera sensor during acquisition according to

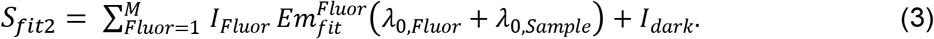

The analysis did not benefit from including a time-varying fit shift of the emission spectra for each camera frame, which would have accounted for alignment shifts during the 30-sec acquisition if present. The sample-specific sub-pixel calibration shifts reduced the unexpected positive and negative cross-correlation amplitudes. Through the above, multi-step method, many variables are set precisely to well-resolved spectra such that the potentially noisy experimental spectra from a single camera frame with highly discrepant fluorophore concentrations could be fit with minimal fitting unknowns; only the intensity of each fluorophore unknown and fit to each camera frame in the experimental data.

With the shifted continuous fluorophore emission spectra identified, the expected intensity from each fluorophore on each camera pixel was determined and linear least squared fitting was performed to each camera frame to provide the intensity of each fluorophore at each moment in time. Linear least squared fitting provides 100x improved computational efficiency and indistinguishable auto- and cross-correlation results from a nonlinear least square fitting of the continuous sum of 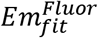 to the experimental spectra.

### 2.4 Data analysis

The autocorrelation (*G*^*Auto*^) of each intensity versus time signal revealed the diffusive behavior via the correlated intensity versus lag time (*τ*) of a single species according to

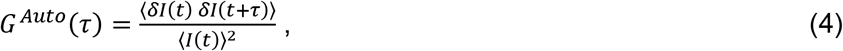

where 〈*I*(*t*)〉 represents the time average intensity. In FCCS, two emission intensities versus time are analyzed via cross-correlation (*G*^*Cross*^) according to

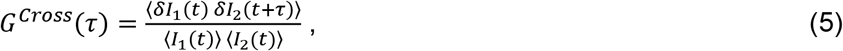

The auto and cross-correlation were fit to the expected correlation decay of homogeneous 2D or 3D Brownian diffusion (Eq. 6 or 7) to extract the correlation amplitude (*G_0_*) and *τ*_*D*_ (36) according to

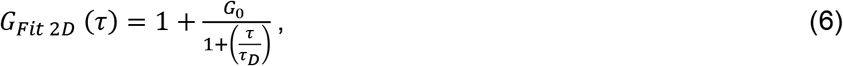

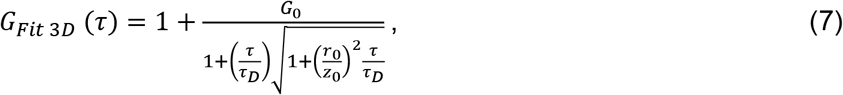

Eq. 7 incorporates the radial (*r_0_* = 0.61*λ*/NA) and axial (*z_0_* = 2n*λ*/NA^2^) distances over which the intensity decay by 1/e^2^, the immersion oil index of refraction (*n* = 1.51) and the numeric aperture of the objective (*NA* = 1.30) (37). The acquired intensity versus time traces were checked for photobleaching and significant stage drift before fitting to Eqs. 6 or 7.

*G_0_* for autocorrelations 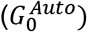 reports the number of diffusers in the detection volume (*N* ∝ *1/G_0_*). Interpretation of the magnitudes of cross-correlations 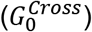 was performed by calculating the fraction of correlated (*F_c_*), according to

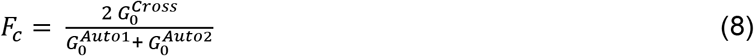

## 3. Results

### 3.1 Comparison of sCMOS vs EMCCD cameras

The read noise of sCMOS is low compared to EMCCD, but the dark counts were lower, and the quantum efficiency was greater for the EMCCD. Both the sCMOS and EMCCD cameras worked well to detect the transit of bright diffusers that had multiple fluorescent molecules each (*i.e*., the nanobeads and the LUVs). However, the EMCCD’s provided superior signal to noise for the single-fluorophore diffusers (*i.e*., the single proteins or lipids). Both cameras enabled sufficient hardware and software for this method, but with our need for a system that is suited for many sample types and the challenges of switching between cameras, we chose to primarily use EMCCD.

### 3.2 Induced oligomerization of nanobeads

The diffusion and oligomerization of the fluorescent nanobeads or LUVs were measured in aqueous suspensions. Four distinct types of polystyrene nanobeads were measured with peak excitation wavelengths of 450, 515, 561, or 635 nm (B_450_, B_515_, B_561_, or B_635_). Three different types of LUVs were labeled with TopFluor-PC (TF), DPPE-Texas Red (TR), or DOPE-Cy5 (CY5) with peak excitation wavelengths of 515, 561, or 647 nm (L_515_, L_561_, or L_647_). Nanobeads or LUVs were mixed, and their fluorescence emission intensities were collected for 30 sec at >1.7 kHz. Intensities versus time for each channel showed many small and uncorrelated events that reflect independent diffusion of each species (Figs. 3A and S4A). The average of 27 sequential measurements of autocorrelations for four channels were calculated and fit to yield *τ_D_* for each bead type (Fig. 4B), which were in accordance with the expected values for the varying size beads (Table 1). In ultrapure water, the cross-correlations between the nanobeads were negligibly small or negative (Figs. 3C, 4C).

**Table 1:** Comparison of beads’ diffusion, diameter, and oligomer fraction.

| Bead Type | Diameter from manufacturer (nm) | D<sub>Theory</sub> (μm<sup>2</sup>/sec) | D<sub>Exp</sub> (μm<sup>2</sup>/sec) |  | Estimated diffuser diameter (nm) |  | Oligomer Fraction (%) |
| --- | --- | --- | --- | --- | --- | --- | --- |
|  |  |  | Water | PBS | Water | PBS | PBS |
| B<sub>450</sub> | 100 | 4.39 | 4.45 ± 0.06 | 0.71 ± 0.05 | 98.7 ± 1.2 | 618.7 ± 40 | 99.9 |
| B<sub>515</sub> | 100 | 4.39 | 2.86 ± 0.03 | 0.92 ± 0.07 | 153.9 ± 1.4 | 477.5 ± 30 | 99.7 |
| B<sub>561</sub> | 100 | 4.39 | 3.44 ± 0.02 | 0.80 ± 0.06 | 127.7 ± 0.6 | 547.7 ± 40 | 94.4 |
| B<sub>635</sub> | 40 | 10.98 | 7.09 ± 0.06 | 1.00 ± 0.06 | 61.9 ± 0.5 | 439.3 ± 20 | 97.5 |

**Figure 3:**
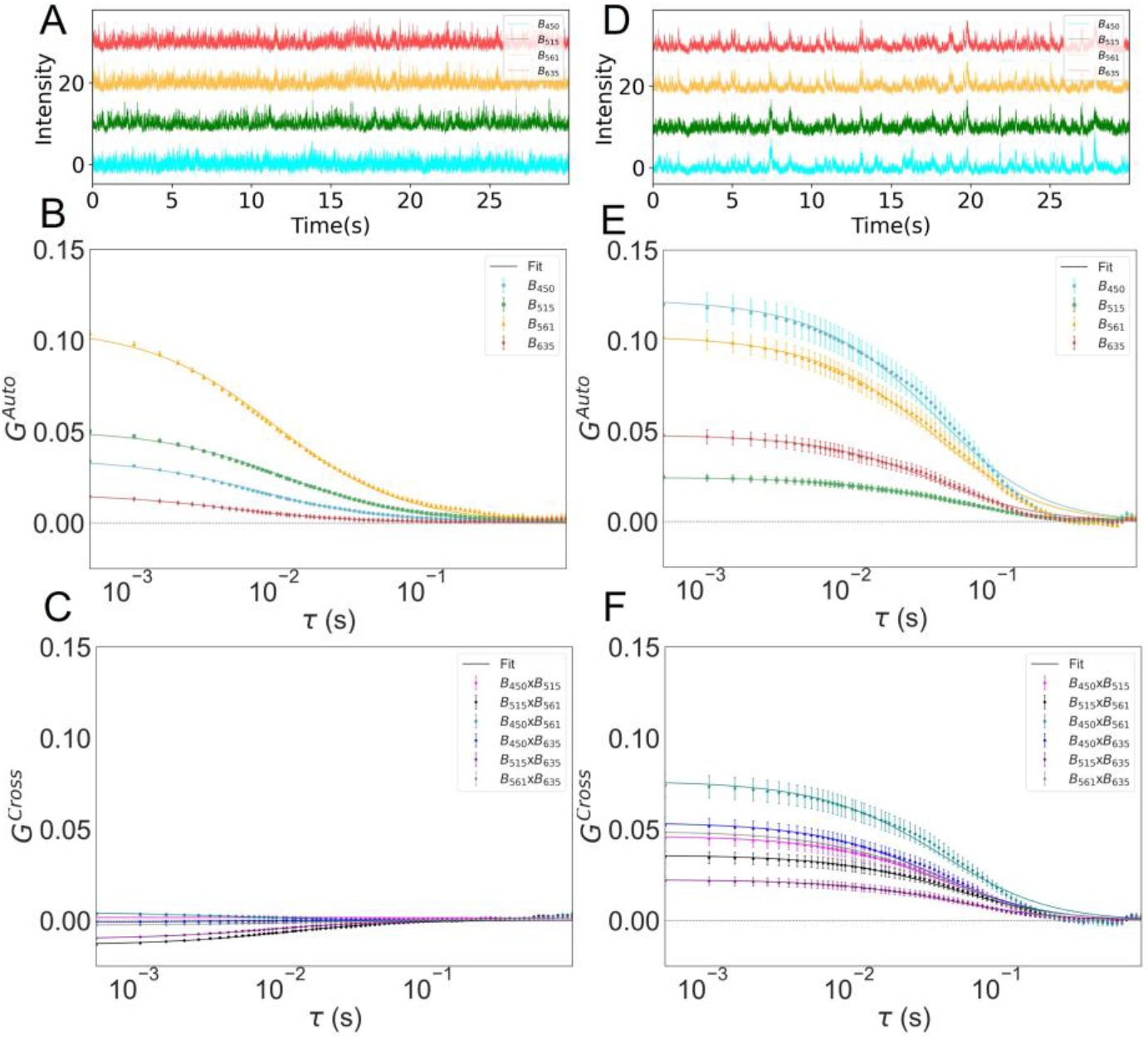
FCCS on four spectrally overlapping nanobeads. (A) Fluctuations in intensity versus time for four nanobeads (B_450_, B_515,_ B_561,_ B_635_) show many small uncorrelated peaks that indicate the nanobeads are diffusing independently. The results were analyzed with (B) auto and (C) cross-correlations. (B) The autocorrelations were well resolved and (C) the cross-correlations were small or negative. The addition of PBS induced aggregation of nanobeads. (D) The intensity versus time for the four-color channels showed many events with correlated peaks because the nanobeads formed clusters and were diffusing together. PBS yielded both the (E) auto- and (F) cross-correlations increased to display increased amplitudes and dwell times. (B,C,E,F) Symbols and error bars represent the mean and standard error of the mean for >20 sequential measurements while the lines represent the fits from Eq. 7.

**Figure 4:**
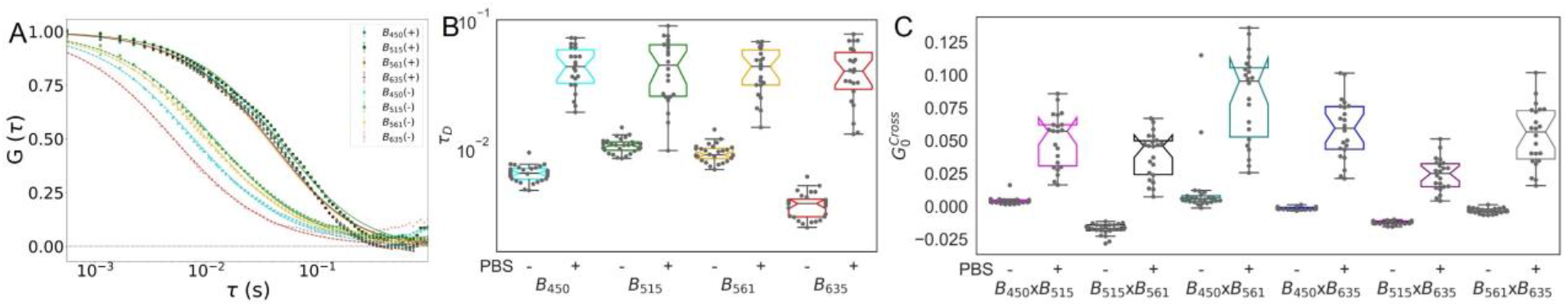
(A) The normalized autocorrelations in water and PBS for the four nanobeads simultaneously observed. A minus sign (−) indicates the absence of PBS and the plus sign (+) indicates the presence of PBS. (B) The autocorrelations displayed slower diffusion upon the addition of PBS and (C) the cross-correlations displayed larger amplitudes upon the addition of PBS, which demonstrates that heterooligomers had been formed.

Upon increasing the salinity of the solution by the addition of PBS, the nanobeads randomly bound together as the salts encouraged oligomerization. The four intensities versus time showed many correlated peaks confirming the formation of large, multicolored nanobead clusters (Fig. 3D). Upon oligomerization, the autocorrelations showed longer decay times as the diffusion slowed (Figs. 3E, 4B). This is best demonstrated by comparing normalized autocorrelations with water or PBS; increasing salinity caused the beads to clump and diffuse slower (Fig. 4A). Additionally, the cross-correlations displayed significantly increased G_0^Cross and FC as the binding increased. (Fig. 3F, 4C).

### 3.3 Oligomer density of nanobeads

The composition of the oligomers was examined by recognizing that the autocorrelations included information about both the monomeric and oligomerized diffusers within a sample. The autocorrelations in water provide the diffusion dynamics of only the monomers. The cross-correlations in PBS provide the diffusion dynamics of only the oligomers. The autocorrelations in PBS include the dynamics of both the monomers and the oligomers as a reflection of the polydispersity of the sample. Accordingly, we fit the autocorrelation in PBS to be a linear sum of the autocorrelation in water and the cross-correlation in PBS such that the relative amplitudes of the components reflected the fraction of the diffusers that were monomers vs oligomers, according to

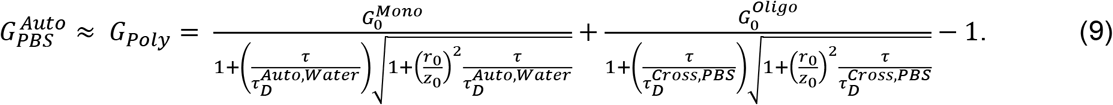

The resulting 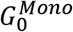 and 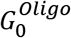 were used to calculate the fraction oligomerized according to

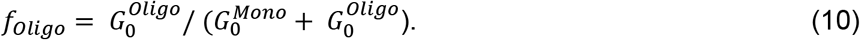

and listed within Table 1. We observed all four nanobeads displayed *f*_*Oligo*_ above 94%, with B_450_ having *f*_*Oligo*_ equal to 99.9%. Analysis of the bead monomer and oligomer diffusion coefficients, the oligomer diameters were estimated to be 520 ± 90 nm, which was 6x larger than that of the monomeric beads. This indicates that over 200 nanobeads joined into a single oligomer (Table 1). Similarly, we observed all three LUVs displayed *f*_*Oligo*_ above 77% (Table S2). The Stokes-Einstein relationship was used to connect the diffuser diameter to diffusion coefficient while assuming a solution viscosity of 1 cP and temperature of 20 °C.

### 3.4 Induced oligomerization of lipid vesicles

Three-color FCCS was also performed on LUVs that incorporated three highly overlapping fluorescent lipids, as described above. The fitting of continuous functions to the individual fluorophore’s spectra, shifting the fits to accommodate optical misalignments, and the linear least squared fitting processes described above effectively separated the combined spectra for extraction of each fluorophore’s intensity vs time (Fig. S3). Our observations were like the nanobeads in that the LUVs were diffusing independently in water (Fig. S4A, B) whereas started forming homo- and heterooligomers showing strong cross-correlations were induced. With the LUVs, heterooligomerization was induced by the addition of BSA at 30 g/L to the buffer (Fig. S4C, D).

### 3.5 Membrane-bound protein diffusion on lipid bilayers

To evaluate the sensitivity of our system to single, membrane-bound molecules, we examined membrane-bound fluorescent proteins of cholera toxins (CTxB). CTxB are soluble proteins that bind to the glycolipid GM1 incorporated into phospholipid membranes. The CTxB that was labeled with the synthetic, bright AlexaFluor (AF) dyes were successfully perform with either the sCMOS or EMCCD camera. An SLB sample with both CTxB-AF488 and CTxB-AF647 bound displayed independent diffusion of the two populations of CTxB and no cross-correlation (Fig. 5A, B).

**Figure 5:**
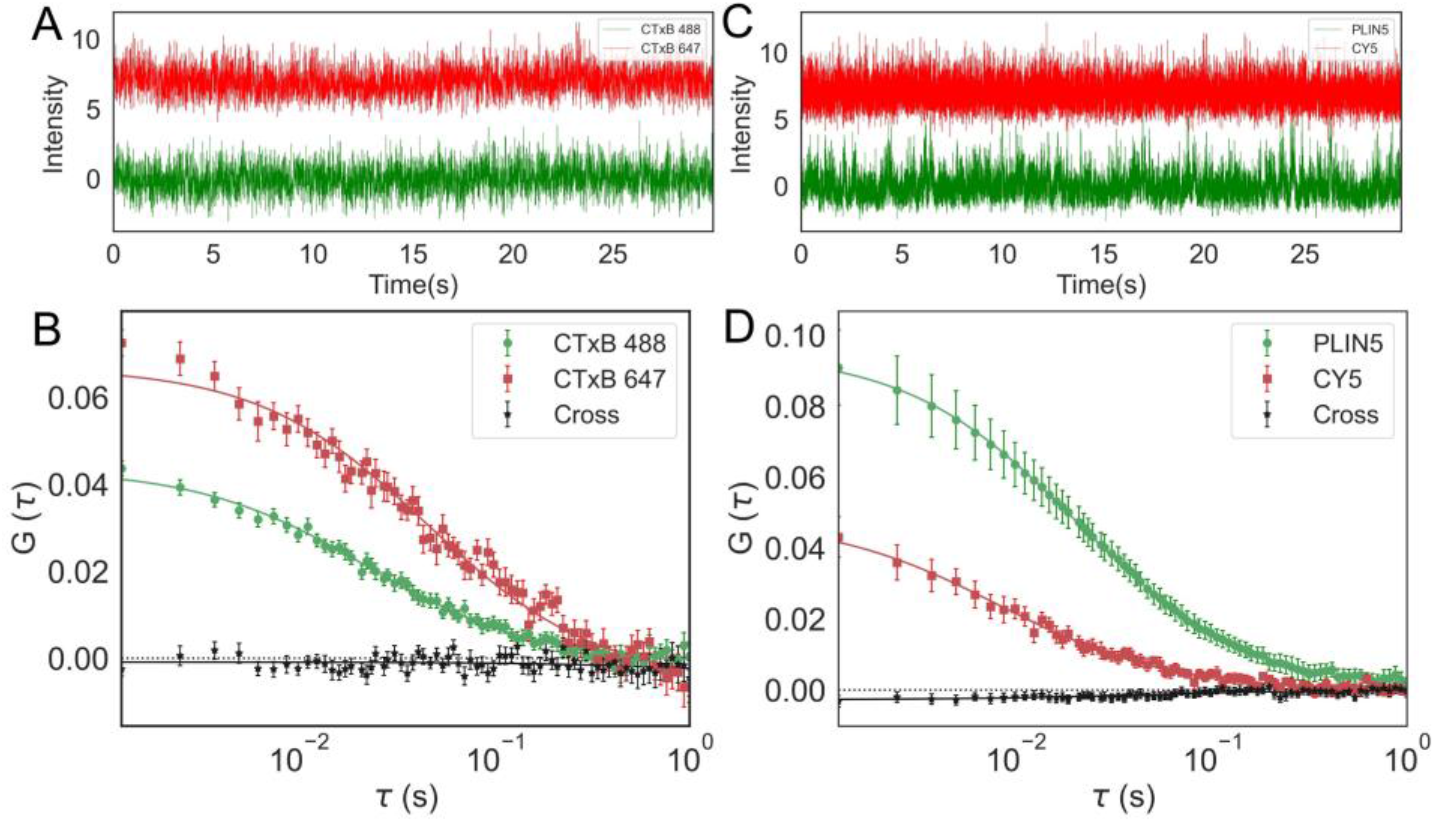
Protein-protein and protein-lipid diffusion on phospholipid bilayer and monolayer were detected by our FCCS system. (A) The intensity versus time curves of two proteins CTxB-AF488 and CTxB-AF647 were analyzed to yield (B) two autocorrelations and cross-correlation of these bilayer-bound proteins. No apparent association between the two distinct populations of CTxB was observed. (C) The intensity versus time PLIN5-YFP and DOPE-CY5 were analyzed to yield (D) two autocorrelations and a cross-correlation of the monolayer-associated molecules. PLIN5-YFP and DOPE-Cy5 diffused independently on surface of lipid monolayer. Symbols and error bars represent the mean and standard error of the mean for >25 sequential measurements while the lines represent the fits from Eq. 6.

### 3.6 Independent protein and phospholipid diffusion on lipid monolayer

To evaluate the capability of our system to measure the diffusion on the phospholipid monolayer surface of an aLD, we examined the diffusion of a fluorescent phospholipid (DOPE-Cy5) and a purified lipid droplet scaffolding protein labeled with a yellow fluorescent protein (PLIN5-YFP). FCCS was performed on the “south pole” of the aLDs and revealed no cross-correlation between DOPE-Cy5 and PLIN5-YFP. The autocorrelations demonstrated that DOPC-Cy5 was diffusing 2x faster than PLIN5-YFP, and the cross-correlations were negligible (Fig. 5C, D).

## 4. Discussion

With only one laser source and one detector, our custom FCCS reveals four distinct color channels for a novel experimental routine. No cross-correlations were observed for independent diffusers despite highly overlapping spectra. FCCS data are important for providing information on the mobility, concentration, and diffusion dynamics through the analysis of autocorrelations. Amplitudes of autocorrelations are inversely proportional to number of diffusers in observation volume. Also, *τ_D_* can be quantified in each of auto- and cross-correlations that gives residence time of diffusers in the detection volume. In addition, the multicolored clusters can be evaluated by analyzing cross-correlation amplitudes.

All the nanobeads were moving independently to each other in ultrapure water; cross-correlations did not give positive amplitudes. Small and negative cross-correlation magnitudes between the beads in ultrapure water were analyzed for potential excluded volume effects, spectral bleed through, physical bead sizes, and surface chemistry of the beads. An excluded volume effect may have contributed to the negative cross-correlations (*i.e*., 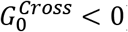). The presence of one 100-nm diameter bead within the 300-nm wide observation volume likely reduced the probability of other beads in the observation volume simultaneously, thus the negative correlation between beads (2, 38). Comparing beads of differing diameter did not show a clear correlation between bead sizes and 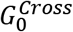 (Fig. 6).

**Figure 6:**
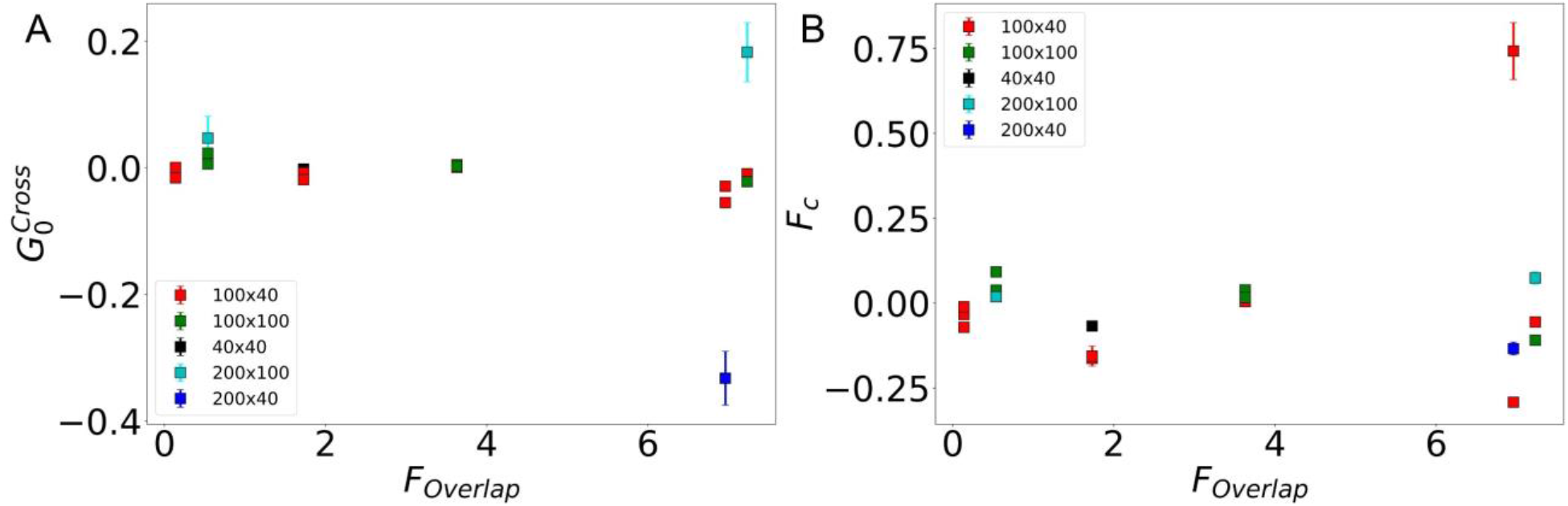
Fractional overlapping of the wavelengths versus bead parameters. (A) *F_Overlap_* versus 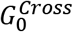 for combination of bead sizes (B) Fraction correlated (F_c_) for varying bead sizes plotted against *F_Overlap_*.

The induction of nanobead oligomerization with addition of PBS shows increase in *G_0_* in all six pairs of cross-correlations (Fig. 4C). The larger amplitude means fewer independent diffusers, consistent with the individual diffusers clumping together to form multicolored clusters. *G_0_* for autocorrelations also increased, with the exception of B_515_ (Fig. S2). Upon aggregation, *τ*_*D*_ increased 1.8 ± 0.7 ms, with B_635_ having maximum 3.00 ± 0.03 ms increase compared to ultrapure water *τ_D_* for same beads (Fig. 4B).

To examine the contribution of spectral overlap to the observed cross-correlation, we computed fractional spectral overlap (*F_Overlap_*) between each pair of beads according to

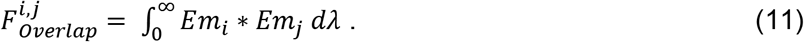

*Em_i_* is the normalized emission spectrum for bead *i*. *F_Overlap_* ranged from 0.14% to 7.23% while 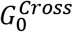 was between −0.35 to 0.18. There was no correlation between *F_Overlap_* and 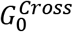 for any of the nanobeads (Fig. 6A). As expected far wavelengths had less spectral overlapping than the closer ones, but they had no greater 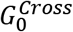 or *F_C_* for bead of varying physical sizes (Fig. 6B). Further the anti-cross-correlation which might be the cause of bleed through between the channels, but we do observe the negative cross-correlation for channels B_450_xB_635_ which is 450 nm and 635 nm emission channels that has negligible (*F_Overlap_* = 0.14%) bleed through between them. So, Bleed through cannot lead to anti-cross-correlation.

Moreover, to access surface chemistry of the beads, the beads’ surfaces were coated either with uncharged polystyrene or were negatively charged carboxylate coated (Table S1). None of the combinations of bead surface chemistries showed any significant trends for 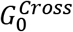. We found no trends for the observed negative cross-correlation with dependences on the bead color, spectral overlap, size, or surface chemistry.

To investigate the possibility for data analysis artifacts being the source of anti-cross-correlation, we developed the computational routine where one intensity is iteratively allowed to fit to sum of calibrations and rest of the other intensities were kept constant. This method provided a new intensity versus time curve for each of the four-color channels, but extracted correlations gave false positive cross-correlations. It seems like fluctuations in the total laser power may have resulted in this false positive cross-correlation. Also, the background intensity (*I_dark_*) for the correlation analysis upon examination showed that there was no interdependency between correlations and *I_dark_*. Further, fitting for fluorophore spectra that were not present in the sample yielded the expected near zero intensity for these intensities and no changes to the cross-correlations for the present fluorophores.

The four intensities present for four beads allowed us to examine if all beads contributed similarly to the oligomerization. Three and four intensities were correlated to acquire two and three lag times between them, respectively (Eqs. S1, S2). The 2D plot of cross-correlation matrix showed *τ*_1_ and *τ*_2_ are independent and perpendicular to each other for nanobeads in PBS (Fig. S5)(39). Three nanobeads were chosen at a time for this analysis to assess association and dissociation between macromolecules of interest and assess if the interaction of any two particular bead types were key to the larger, multi-color aggregation. We did not observe the key large multicolored aggregation between the nanobeads or in LUVs (Fig. S6) via double or triple cross-correlation analysis, but this quantification routine can be useful for the future experiments that needs to examine multi-color protein aggregation and complex assembly.

In sum, our FCCS has a wide application on variety of samples including live cells, lipid monolayers, and bilayers. The limitation of the system is misalignment of either pinhole or camera and drift in focus, all of which can lead to noisy autocorrelations. But which can tackle easily following strict experimental and statistical routines for focused sample between excitation and emission channel.

## 5. Conclusions

In this study, we developed a multi-color FCCS system that demonstrated interaction and diffusion dynamics of four spectrally overlapping fluorescent probes. Our economical FCCS uses multiple laser channels in the excitation path that were created from single supercontinuum laser (400-700 nm) and a dispersive prism that gave the emission spectra of multiple chromatically overlapping fluorophores on to a single detector. Measurements of diffusion time and fluorophore density were successfully performed on polystyrene fluorescent nanobeads. Also, on the same nanobeads system FCCS enabled us to observe induced aggregation and clustered interaction caused due to addition of PBS. Measurement of diffusion coefficient clearly showed long decay time and slower diffusion in presence of PBS due to formation of multicolored oligomers. Due to the adaptiveness of accommodating up to four molecules at a time, our FCCS is conventional to use with fluorescent proteins and lipids in biological model and live systems. Moreover, the ability to calculate full triple cross-correlation decay, the size of the three-color species can be possible to determine via FCCS. The advantage of the system will be investigation of diffusion simultaneously with same experimental conditions on multiple fluorophores that will also save multiple sample preparation and experiment time. Future experiments will include up to four lipolysis associated proteins and phospholipids, that will be examined on the surface of monolayers. This approach will be used to investigate triple cross-correlation decay and can identify complex assembly and disassembly kinetics.

## Acknowledgements

This material is based upon work supported by the National Science Foundation under Grant No. DMR1652316. Research reported in this publication was supported by the National Institute of Diabetes and Digestive and Kidney Diseases of the National Institutes of Health under award number R01DK076629. We are grateful to Richard J. Barber for financial support. We thank Matthew Sanders and James Granneman for protein purification and supply. We also thank Susheel Pangeni, Abir Kabbani, Xinxin Woodward and Nasser Junedi for valuable discussions.

## Supplemental Methods

### Protein purification

Baculovirus for protein expression were prepared using the Bac-to-Bac expression system (Invitrogen). pFastBac1 constructs for PLIN5-YFP-His tag (all mouse protein) was made using standard molecular biological methods and was confirmed by sequencing. Bacmid DNAs were generated by transformation of DH10Bac E. coli (Invitrogen) with FastBac plasmids following the manufacturer’s protocol. Initial baculovirus stocks were generated by transfection of Sf9 insect cells with bacmid DNA using Cellfectin II reagent (Invitrogen) according to the manufacturer’s protocol, then were amplified by infection of Sf9 cells with the initial baculoviral stocks.

Amplified baculoviral stocks were used to infect High Five insect cells and cells were collected 48 hrs after start of infection when cells were ~80% viable. High Five cells expressing proteins were lysed by sonication in IB containing 20 ug/mL leupeptin and pepstatin A. For PLIN5-YFP-His tag preparations, 0.5% FOS-CHOLINE-12 detergent (Anatrace) was added to sonicated cell extracts prior to centrifugation at 10,000 g. Supernatants were then incubated with TALON Cobalt beads (Takara) for two hours at room temperature, then proteins were eluted from beads using 50 mM sodium phosphate buffer pH 7.4 containing 150 mM imidazole. Imidazole was removed from proteins by successive rounds of concentration and redilution with IB using Centricon centrifugal filter units of the appropriate molecular weight cutoff. Protein concentration was determined by BCA (Pierce), then protein purity was determined from Coomassie-stained PAGE gels.

### Triple cross-correlation data analysis

For the multi-color oligomerization with more than two spectrally resolved molecules, three or four intensity versus time signals may be considered in a single cross-correlation analysis with two or three independent lag times, respectively,(39–42) according to

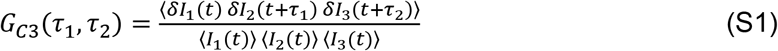

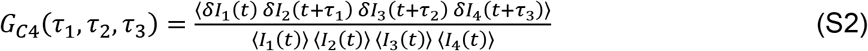

---

**Figure S1:**
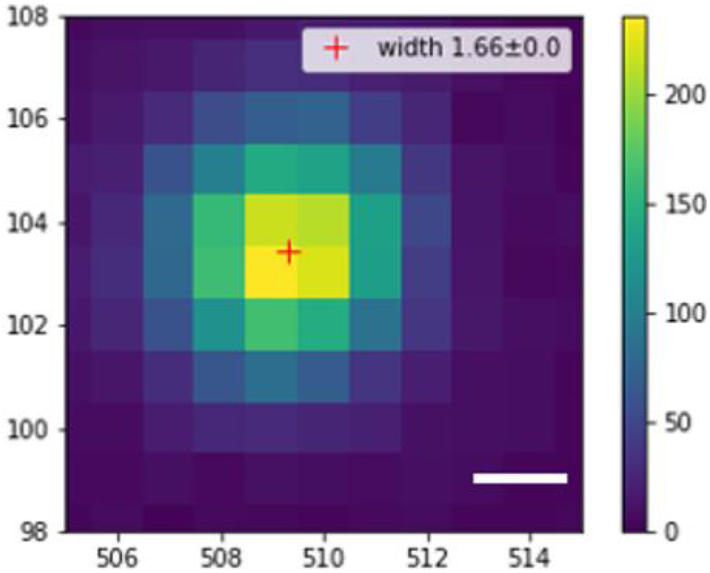
Laser spot size for *λ*_*ex*_ = 561 nm. Scalebar = 0.12 μm.

**Figure S2:**
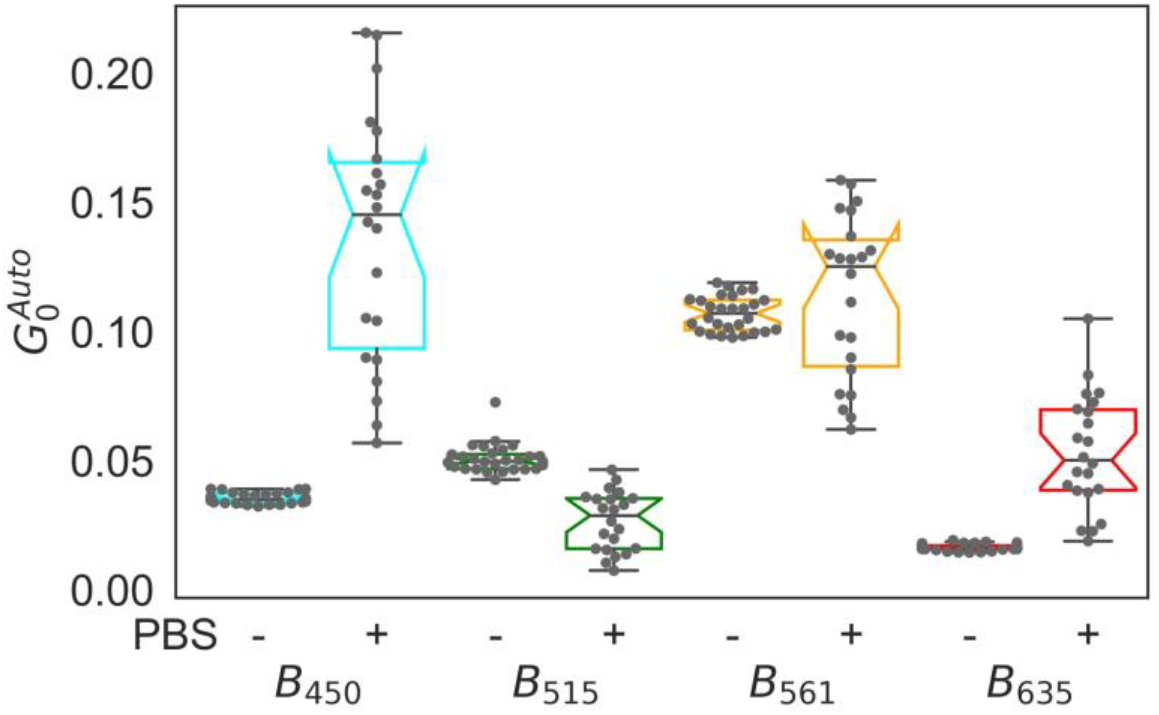
Addition of PBS induced nanobead oligomers. Comparing 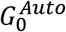 with and without PBS indicates B_450_ and B_635_ have induced homo-oligomers.

**Table S1:** Details of the nanobeads used in these experiments.

| Beads | Size (nm) | Color | Coating | Surface charge | Manufacturer | Lot number |
| --- | --- | --- | --- | --- | --- | --- |
| $B_{450}$ | 100 | Blue | Polystyrene | neutral | Thermo Fisher | 184539 |
| $B_{515}$ | 100 | Yellow | Carboxylate | negative | Life Technologies | 1588588 |
| $B_{561}$ | 100 | Red | Polystyrene | neutral | Thermo Fisher | 180315 |
| $B_{635}$ | 40 | Dark red | Carboxylate | negative | Life Technologies | 1437595 |

**Figure S3:**
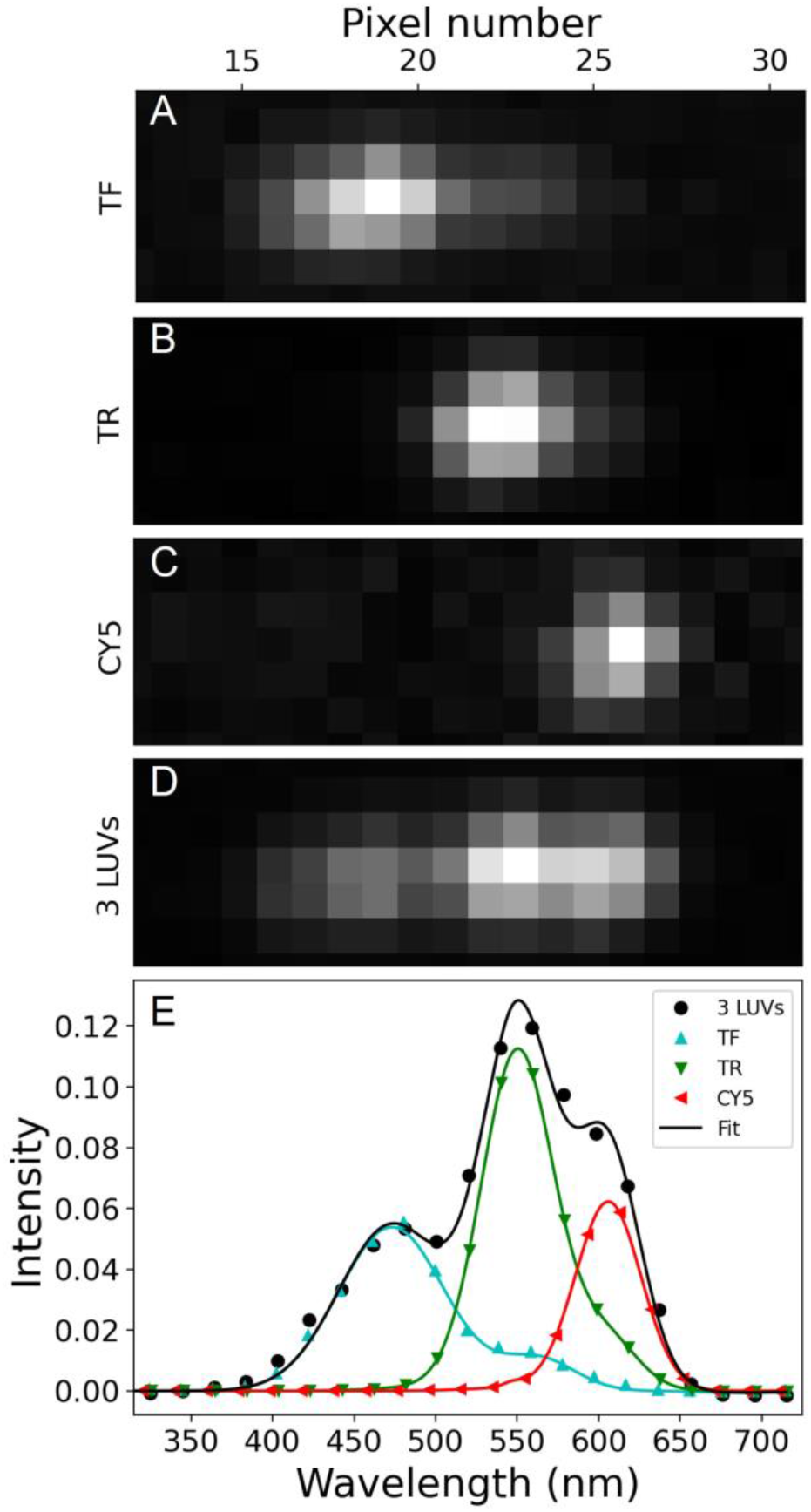
The emission spectra of the LUVs on the camera sensor. (A-C) Single LUVs TF, TR and CY5 of excitation wavelengths 515, 561 and 635 nm, respectively, were spectrally separated and collected on the cropped ROI. Each column of pixels on the camera is associated with an emission wavelength. (D) Samples with all three LUVs show the distinct, but highly overlapping emission spectra. (E) Control samples with only one color of LUV present were used to identify the emission spectrum of each LUV type (*colored symbols and fits)*. With three LUVs simultaneously present, the spectrum was fit to reveal the emission intensity of each LUV type for each camera frame (*black symbols and fit)*.

**Figure S4:**
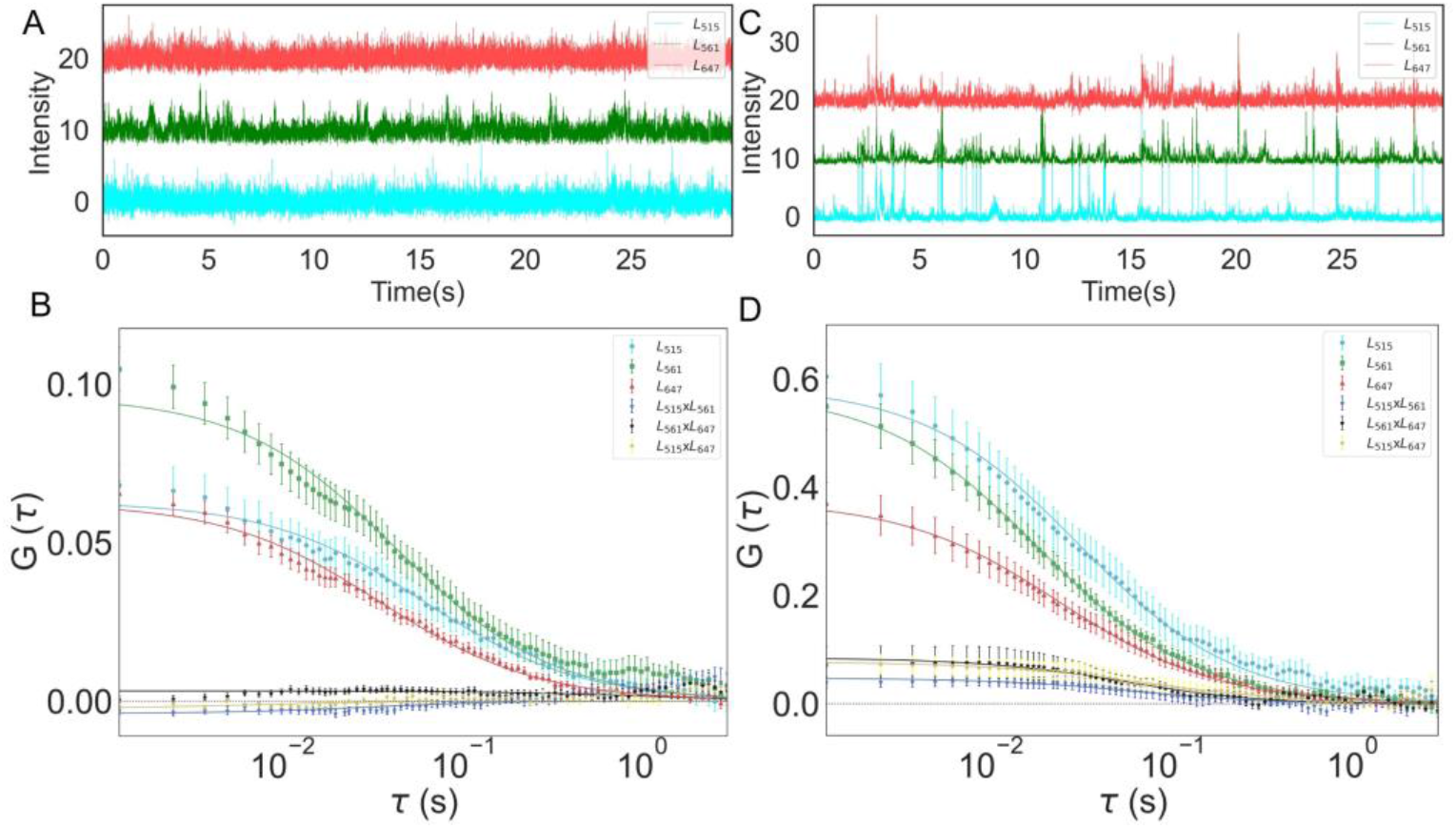
LUVs form homo-and heterooligomers with addition of BSA. (A) Intensity versus time for three LUVs (TF, TR, CY5) in water shows many small uncorrelated peaks that confirm LUVs are diffusing independently. (B) G versus *τ* reports auto correlations were significant whereas cross-correlations were unclear. (C) Addition of BSA induces aggregation of LUVs, intensity versus time for three color channel shows many events of correlated peaks. (B) G versus *τ* reports both auto and cross-correlations were significant, induced aggregation cause increase in auto- and cross-correlations. Symbols and error bars represent the mean and standard error of the mean for >7 sequential measurements while the lines represent the fits from Eq. 7.

**Table S2:**
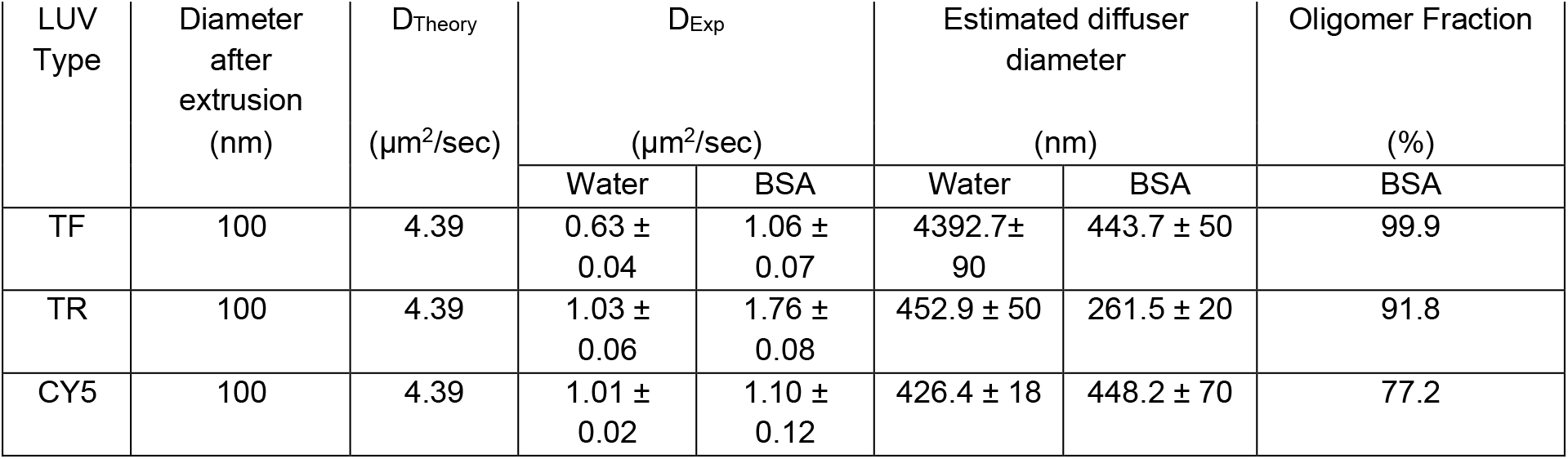
Comparison of LUV diffusion, diameter, and oligomer fraction.

| LUV Type | Diameter after extrusion (nm) | $D_{\text{Theory}}$ ($\mu\text{m}^2/\text{sec}$) | $D_{\text{Exp}}$ ($\mu\text{m}^2/\text{sec}$) | | Estimated diffuser diameter (nm) | | Oligomer Fraction (%) |
| --- | --- | --- | --- | --- | --- | --- | --- |
|  |  |  | Water | BSA | Water | BSA | BSA |
| TF | 100 | 4.39 | $0.63 \pm 0.04$ | $1.06 \pm 0.07$ | $4392.7 \pm 90$ | $443.7 \pm 50$ | 99.9 |
| TR | 100 | 4.39 | $1.03 \pm 0.06$ | $1.76 \pm 0.08$ | $452.9 \pm 50$ | $261.5 \pm 20$ | 91.8 |
| CY5 | 100 | 4.39 | $1.01 \pm 0.02$ | $1.10 \pm 0.12$ | $426.4 \pm 18$ | $448.2 \pm 70$ | 77.2 |

**Figure S5:**
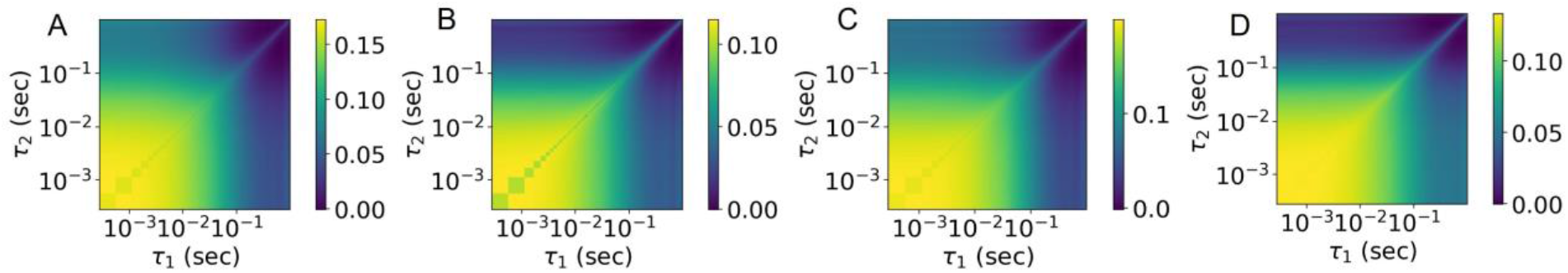
Contour plots of three-color cross-correlation decay for nanobeads. The combination of three beads at a time out of four nanobeads was examined to see macromolecular correlations following Eq. S1.

**Figure S6:**
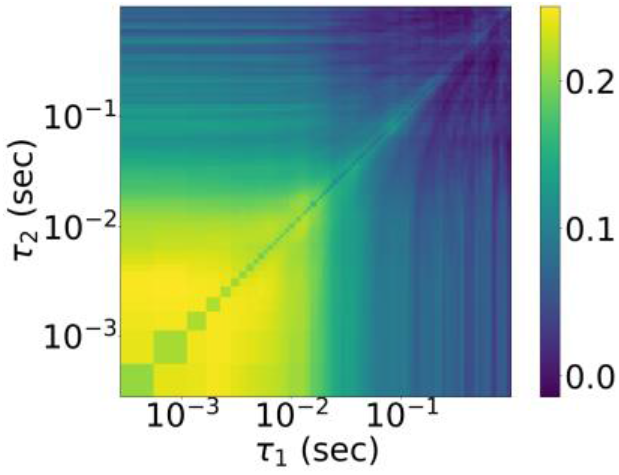
Contour plot of three-color cross-correlation decay for LUVs (TF, TR, CY5).

